# Likelihood-based interactive local docking into cryo-EM maps in ChimeraX

**DOI:** 10.1101/2024.05.16.594509

**Authors:** Randy J. Read, Eric F. Pettersen, Airlie J. McCoy, Tristan I. Croll, Thomas C. Terwilliger, Billy K. Poon, Elaine C. Meng, Dorothee Liebschner, Paul D. Adams

## Abstract

The interpretation of cryo-EM maps often includes the docking of known or predicted structures of the components, which is particularly useful when the map resolution is worse than 4 Å. Although it can be effective to search the entire map to find the best placement of a component, the process can be slow when the maps are large. However, frequently there is a well-founded hypothesis about where particular components are located. In such cases, a local search using a map subvolume will be much faster because the search volume is smaller, and more sensitive because optimizing the search volume for the rotation search step enhances signal-to-noise. A Fourier-space likelihood-based local search approach, based on the previously-published *em_placement* software, has been implemented in the new *emplace_local* program. Tests confirm that the local search approach enhances speed and sensitivity of the computations. An interactive graphical interface in the ChimeraX molecular graphics program provides a convenient way to set up and evaluate docking calculations, particularly in defining the part of the map into which the components should be placed.

**Synopsis:** Likelihood-based cryo-EM docking using our *emplace_local* software is faster and more sensitive than our related software, *em_placement*, when the approximate location of a component is known, and is available conveniently through a plugin to the ChimeraX visualization software.

## 1 Introduction

Over the past decade, improvements in cryo-EM hardware and algorithms have led to an explosion of new maps, many (but not all) at resolutions that permit atomic modelling of proteins and other biological macromolecules. When atomic models of the same or closely-related components are available – either from the Protein Data Bank (PDB) (Berman *et al*., 2007) or from structure prediction algorithms such as AlphaFold (Jumper *et al*., 2021) or RoseTTAFold (Baek *et al*., 2021) – docking techniques can quickly yield an initial model.

Docking is particularly important when the quality of the reconstruction is limited (*e*.*g*. by overall or local resolution poorer than 4 Å), so that an atomic model is difficult to build *ab initio*. In such cases, docking success will depend on making the best possible use of the signal in the data.

For this reason, we have pursued a likelihood-based approach to docking, which accounts for the effects of errors in the cryo-EM reconstruction and in the search model. Likelihood also has the advantage that the likelihood score itself allows one to infer a level of confidence in a docking solution. Specifically, a likelihood score of 60 or more is extremely unlikely to be random so it indicates a correct, or at least partially correct, solution (McCoy *et al*., 2017).

The likelihood target for docking is expressed in Fourier space, similar to the approach of modern likelihood-based cryo-EM reconstruction algorithms such as RELION (Kimanius *et al*., 2021). Previous papers describe the theoretical background (Read *et al*., 2023) and the implementation and testing (Millán *et al*., 2023) of the docking procedure in the *em_placement* program, which searches globally over the entire map to locate the molecule of interest. The test calculations covered a range of map resolutions (1.7-8.5 Å) and model completeness (about 5-50% of the total reconstruction), and most of them succeeded.

However, some of the model placements failed when the signal-to-noise in the search over the whole map was too poor. Here we examine why such docking calculations fail and how some can be rescued if the user has a correct hypothesis for the location of the component. The local search, implemented in *emplace_local*, is fast enough to be run interactively through a graphical interface, and is available in the UCSF ChimeraX framework (Goddard *et al*., 2018).

## 2 Local search approach

Before describing the local search technique, it is useful first to review the methodology used in the *em_placement* full search. The six-dimensional problem of finding the orientation and position of a model to fit the reconstruction can be divided into a sequence of two three-dimensional problems: searching for one or several plausible model orientations (rotation function), followed by searching for positions of models in those plausible orientations (translation function). This general approach is used in crystallographic Molecular Replacement (MR), and the Fourier-space equations and algorithms for cryo-EM docking are closely related to reciprocal-space MR equivalents (Read *et al*., 2023).

For MR and particularly for docking, the limiting factor for success is the rotation search, which has intrinsically lower signal than a translation search (carried out for the correct model orientation). In the case of MR, the only ways to improve signal in the rotation search are to find/generate better models or better data/crystals. However, for docking the availability of phase information creates another opportunity. Consider a case, with a low-quality map and an incomplete model, in which the signal in the rotation search is too weak to detect the orientation of the modelled component in Fourier terms computed from the full map. If a smaller volume of the map containing the target component is used to prepare Fourier terms for the search, the noise from unexplained map features will be reduced and the signal-to-noise for the rotation search will be increased. As long as the cut-out volume contains the target component, the signal-to-noise in the rotation search, assessed using the rotation function expected log-likelihood-gain (eLLG_rot_), is predicted to be inversely proportional to the search volume (Read *et al*., 2023).

The decision making in the global search procedure therefore relies on the signal available for the rotation function (Millán *et al*., 2023). If an analysis over the full reconstruction suggests that there is sufficient signal to expect the correct orientation to be in the list of plausible orientations (eLLG_rot_ > 7.5), a single rotation search is performed using the Fourier terms derived from the full reconstruction. If not, rotation searches are performed over smaller subvolumes, a set of spheres that cover the full reconstruction with sufficient overlap to ensure that the full volume of the target will be enclosed in at least one sphere. The size of the subvolumes is chosen (based on the statistical analysis of the full reconstruction) to be as large as possible to limit the number of separate searches while retaining sufficient signal.

Note that the number of subvolume spheres required to cover the full reconstruction with sufficient overlap grows rapidly if the subvolume radius drops below about 1.15 times the model radius, so there is a practical limit to how small these volumes can be made.

The strategy of performing the rotation search on subvolumes depends on the implicit assumption that the quality of the reconstruction is similar over the whole map, so that the overall signal is distributed equally over all ordered parts of the reconstruction. If the component being sought is in one of the more poorly-ordered parts of the reconstruction, the strategy calculations will be overly optimistic and the signal may be inadequate. Note that, although the statistical analysis of signal and noise is carried out separately for each subvolume, *em_placement* does not currently readjust the sizes of the subvolumes in response.

When there is a reasonable hypothesis for the location of the target component, then the search can be optimized based on the principles above. The search can be restricted to the smallest possible subvolume, thereby improving signal-to-noise compared to using larger subvolumes in a global search. Carrying out a local search allows further simplifications that cannot be applied for a global search. Setting up a search over multiple subvolumes requires some knowledge of which regions of the full map contain ordered parts of the reconstruction, so that time is not wasted carrying out computations on the relatively large volumes for disordered solvent regions. Ordered regions of the map can be deduced by computing local averages of the map variance, but that time can be saved if we assume that a local search is carried out in a relevant region. One slight disadvantage of omitting the ordered volume calculation is that only a rough estimate can be made for the fraction of the signal from the search sphere that is accounted for by the model, and the likelihood scoring is thus less well-calibrated. After a series of test calculations, we have chosen by default to estimate the ordered volume in the search sphere as 1.5 times the volume of the search model (meaning that the model is assumed to account for 2/3 of the signal). The ordered volume calculation can still be invoked if half-maps are available, but test cases including those discussed below suggest that the more precise score calibration this allows is not essential for success.

Since 2022, the deposition of half-maps (*i*.*e*. maps obtained from semi-independent reconstructions that each use a random selection of half of the particle images) has been mandatory for new entries in the Electron Microscopy Data Bank (EMDB) (Lawson *et al*., 2016). Such half-maps are required for the analysis of signal and noise in the *em_placement* procedure, as well as for calculating the ordered volume. As a result, the global docking approach cannot be applied to legacy EMDB entries lacking half-maps. For the local docking approach, the intrinsically higher signal-to-noise can often compensate for a less optimal calculation, so we pursued an approximation to the analysis of signal and noise when half-maps are not available.

The docking likelihood target (Read *et al*., 2023) depends on a value for each Fourier term, *D*_*obs*_, which summarizes the effect of the variation in Fourier space of both signal power and noise power. *D*_*obs*_ can be thought of as the equivalent of *FSC*_*ref*_ (the Fourier shell correlation expected between the observed and true maps) for a single Fourier term rather than a whole shell in Fourier space. In the absence of directional information from half-maps, we use the nominal resolution limit, *d*_*min*_, to calibrate a very simple approximation, in which we assume that the *D*_*obs*_ values decrease isotropically with resolution to a value of 0.1 at the resolution limit.

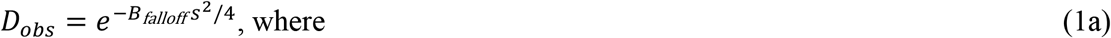

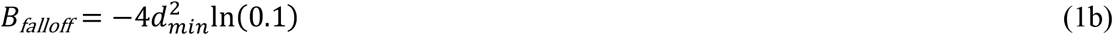

and *s* is the reciprocal of the resolution.

## 3 Implementation of algorithms

### 3.1 Emplace_local

Using functionality previously described for *em_placement*, the *emplace_local* program implements the local search described above. Either a pair of half-maps (preferred) or a single full map must be provided. The search is performed within a sphere, the radius of which is computed automatically by finding the distance to the atom furthest from the center of mass of the search model. The center of the sphere can either be specified as *xyz* coordinates in the 3D space of the map or, if the search model has been placed approximately in the target location, it can be computed from the center of mass of the model. The local resolution of the map in the search volume can be specified (preferred), or a suitable value can be estimated automatically if half-maps have been provided (Millán *et al*., 2023).

Suitable values for local resolution can be estimated with programs such as ResMap (Kucukelbir *et al*., 2014) or *phenix*.*local_resolution* in the Phenix package (Liebschner *et al*., 2019).

### 3.2 Availability in Phenix

The *emplace_local* program is available in the 1.21 release of Phenix and later, and is scheduled to be available in an upcoming release of CCP-EM (Burnley *et al*., 2017). The program can be run from the command-line via the `phenix.voyager.emplace_local` command. With the newer, standard, command-line interface for Phenix programs, the researcher can provide the required information listed above through the parameters available to the program. The new interface also provides a standard command-line flag, `--json`, that will write the output of the program in the standard JavaScript Object Notation (JSON) format that can be easily parsed by other software packages. This approach is used to simplify how the output from this program is interpreted by visualization tools, such as ChimeraX.

### 3.3 ChimeraX plugin

UCSF ChimeraX (Goddard *et al*., 2018; Pettersen *et al*., 2021; Meng *et al*., 2023) is a popular program for the visualization and analysis of biological structural data, ranging from atomic level to cellular or organism level. Recently, the atomic level visualization has been enhanced by the addition of the Isolde plugin (Croll, 2018) for interactive molecular dynamics and building into cryo-EM maps and crystallographic electron density. ChimeraX is now increasingly used for atomic structure display and manipulation and has extensive capabilities for handling cryo-EM maps, making it ideal for presenting a user-friendly front end to the *emplace_local* program. ChimeraX installers for Linux, MacOS or Windows can be downloaded from the ChimeraX website: https://www.cgl.ucsf.edu/chimerax/.

ChimeraX’s plugin architecture not only allows plugins to enhance ChimeraX’s capabilities in many ways, but also allows users to browse and easily install desired plugins from within ChimeraX itself. The PhenixUI plugin, which offers access to *emplace_local* (among other Phenix capabilities), adds an entry to ChimeraX’s Tools menu (“Local EM Fitting”) for launching the graphical front end to *emplace_local*, and adds a command equivalent (“phenix emplaceLocal”) that could be useful for scripting operations. The Phenix software package must be installed separately (https://phenix-online.org), and then the Phenix installation needs to be linked to ChimeraX via the “phenix location” command. For convenience, this command can be added to the ChimeraX Startup items.

The Local EM Fitting tool (Fig. 1) allows the user to specify the structure and cryo-EM map (or preferably half-maps) to use, the local resolution of the map (can be estimated automatically from half maps), and where to initially center the search sphere. If interactive adjustment of the search sphere is enabled in the dialog, then choosing the OK (or Apply) button will show the search sphere (illustrated in section 4.1.3) and allow its position to be interactively adjusted with the mouse. Clicking OK on a secondary dialog will then run an appropriate “phenix emplaceLocal” command which will in turn run the Phenix *emplace_local* program. In addition, the user is offered an option to detect any symmetry that has been imposed on the reconstruction (using the “*measure symmetry*” function in ChimeraX) and to expand the top placed model over that symmetry (“*symmetry*” function).

**Figure 1.**
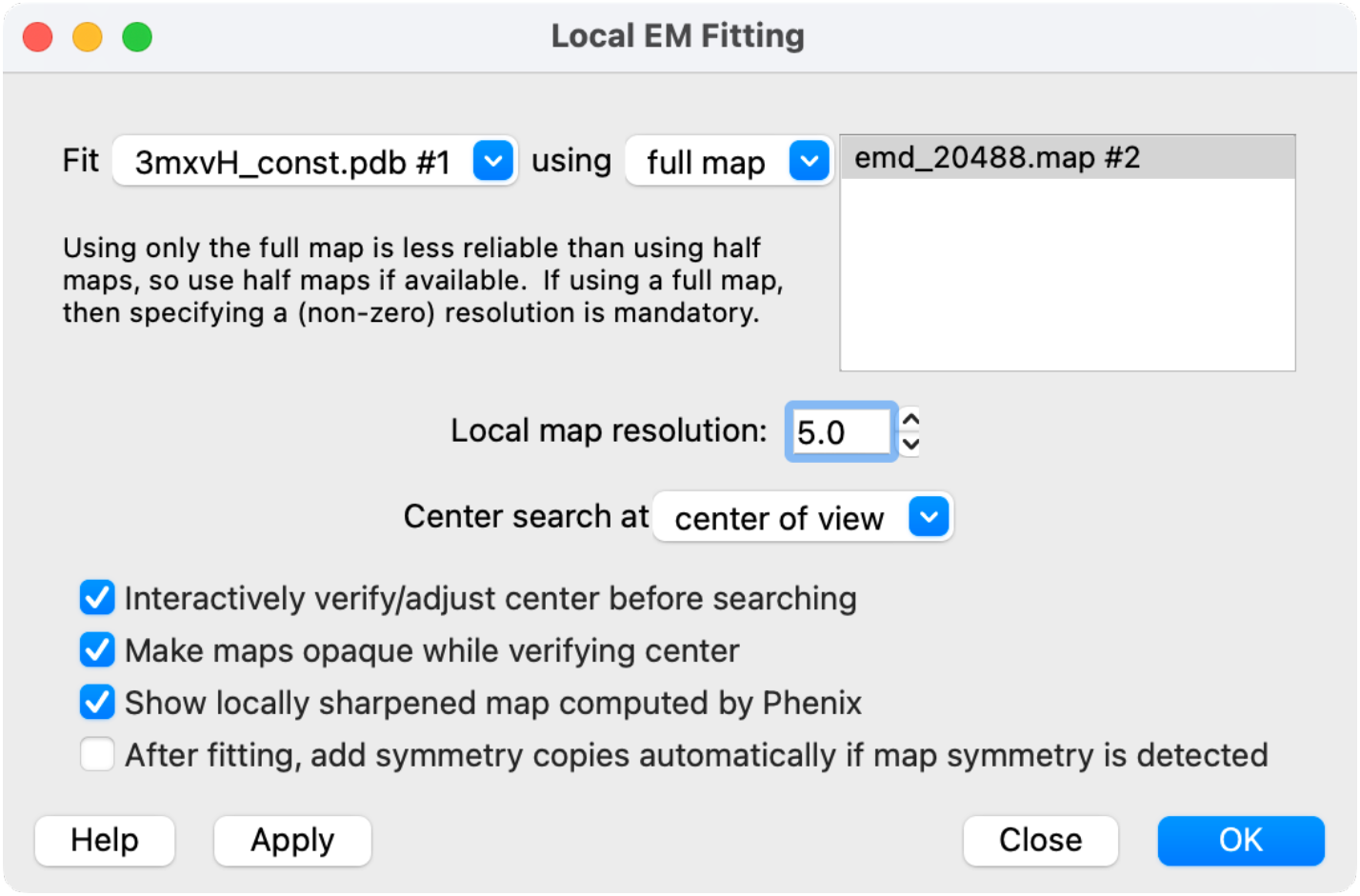
ChimeraX interface to Local EM Fitting tool, illustrating a case discussed in section 4.2.3 below where only a single map is provided and the local resolution is specified.

At the end of the run, the search model is moved to the position found by *emplace_local*, and a “locally sharpened” version of the map is opened. This map is computed from the Fourier coefficients 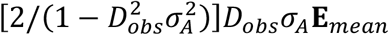 arising from the local map analysis (Millán *et al*., 2023). It provides some visual intuition about the likelihood score, which is roughly proportional to the correlation coefficient between this map and a sharpened map computed from the docked model; even more approximately, the likelihood score would be proportional to the sum of the map values at atomic positions in the model.

## 4 Results

### 4.1 Improved sensitivity and speed of local docking

We reported earlier on the results of global docking tests with *em_placement* on 18 different combinations of docking models and target maps (Millán et al., 2023). By re-examining two of these cases and analyzing a new example, we demonstrate some of the advantages of the local docking search for both speed and effectiveness, when one has a reasonable idea of where a component is located.

#### 4.1.1 Chain L of the *E. coli* respiratory complex I

Some of the most challenging tests were derived from the structure of conformation 2 of the *E. coli* respiratory complex I: PDB entry 7nyu, EMDB entry EMD-12654 (Kolata & Efremov, 2021). Not only is the overall resolution relatively low for *ab initio* modelling at 3.8 Å, but the local resolution varies widely, with map quality dropping dramatically for parts of the membrane domain furthest from the peripheral domain. The poorest local resolution, estimated by the original authors to be in the range 9-11 Å, is for chain L. Map quality improves for the neighbouring chain M and even more for the next neighbour, chain N. Since chains L, M and N are homologous, with about 26% pairwise sequence identities, it is possible to get non-random but suboptimal mismatched placements of the models for these chains.

The global search with chain L of PDB entry 3rko (Efremov & Sazanov, 2011) as a model for the corresponding chain of entry 7nyu failed to find the correct placement; instead, two incorrect placements were found superimposed on chains M and N. An attempt to dock chain M of 3rko succeeded, but the search also yielded an additional incorrect placement superimposed on the more well-ordered map region for chain N. We wished to see whether the *emplace_local* program could be used to match the models of chains L, M and N to their correct sites in the reconstruction, in particular chain L, which failed in the global search.

Table 1 shows the results of local searches pairing each of these models with the three map regions, all carried out at the nominal resolution of 3.8 Å. Note that each of the searches took from 25 to 45 seconds (using a Linux workstation with a 3.8 GHz Intel Core i7-9800X CPU with eight cores but running primarily on a single thread), whether carried out from the standalone Phenix program or the ChimeraX plugin. By comparison, the unsuccessful global search for chain L took 616 seconds on the same computer (Millán et al., 2023). In each case, a search with the model matched to the map region succeeded. However, searches for the mismatched model in the poorest, chain L, map region failed. Searches for the mismatched model in the intermediate-quality chain M region detected part of the map sphere coming from the better-ordered chain N region; as a result, the final rigid-body refinements were no longer centered on the original map region (Table 1a). Carrying out the searches at lower resolution might have been expected to reduce the sensitivity to the local map quality.

**Table 1.** Docking results for alternative pairings of density and model for 7nyu_12654.

| (a) |  | Map target region |  |  |
| --- | --- | --- | --- | --- |
|  |  | chain L | chain M | chain N |
| Model | chain L | L | N‡ | N |
|  | chain M | –† | M | N |
|  | chain N | –† | N‡ | N |

† Docking calculation failed. ‡ A mismatched model was placed in the overlapping map region from the better-ordered neighbouring chain, not the original target region.

| (b) |  | Map target region |  |  |
| --- | --- | --- | --- | --- |
|  |  | chain L | chain M | chain N |
| Model | chain L | 49.9 (0.315) | 88.6 (0.272) | 104.6 (0.279) |
|  | chain M | 4.8 (0.245)† | 241.8 (0.371) | 141.5 (0.276) |
|  | chain N | -1.4 (0.213)† | 57.8 (0.240)† | 486.9 (0.467) |

† Rigid-body refinement was required to enforce the desired solution.

However, when the set of searches was repeated using data limited to either 5, 7 or 9 Å, the same calculations failed with one set of exceptions; using data to between 5 and 9 Å resolution, the chain L model could successfully be docked into the chain M map region.

To obtain docking scores for the desired mismatched pairs, rigid-body refinements were carried out in *em_placement*, starting from a superposition on the target chain from PDB entry 7nyu. The results are shown in Table 1b, with the entries flagged for cases where *emplace_local* failed to yield the desired solution. Likelihood is based on the consistency of a model with a set of data, so it can be used to choose among competing hypotheses for the same data (columns in Table 1b) but not among different data sets for the same hypothesis (rows in Table 1b). With that in mind, it is clear that the model of chain N is the most consistent with the data obtained from the chain N region of the cryo-EM maps, even though non-random scores are obtained for the other models. In contrast, the model of chain L gives the highest score for docking in the chain N region of the map, an intermediate score for docking in the chain M region, and the lowest score for docking in its own region. This does not imply that chain L should really be placed in the chain N position, but just arises because the full map has higher signal to noise in the chain N region, enough to overcome deficiencies of the chain L model in the overall score.

Fig. 2a shows the result of docking chain L of PDB entry 3rko into the chain L region of the target map. The poor quality of the fit to the map is in accordance with a value for the LLG that is close to the level required for confidence. Fig. 2b shows the good superposition of the docked model on the deposited structure in PDB entry 7nyu, along with the deposited reconstruction. We note that the initial fit to this map (Kolata & Efremov, 2021) used models derived from PDB entries 3rko (Efremov & Sazanov, 2011) and 4hea (Baradaran *et al*., 2013), both of which show the same relationship among the components of the membrane arm seen in PDB entry 7nyu. In addition, the authors report carrying out focused refinement, though the resulting map was not deposited.

**Figure 2.**
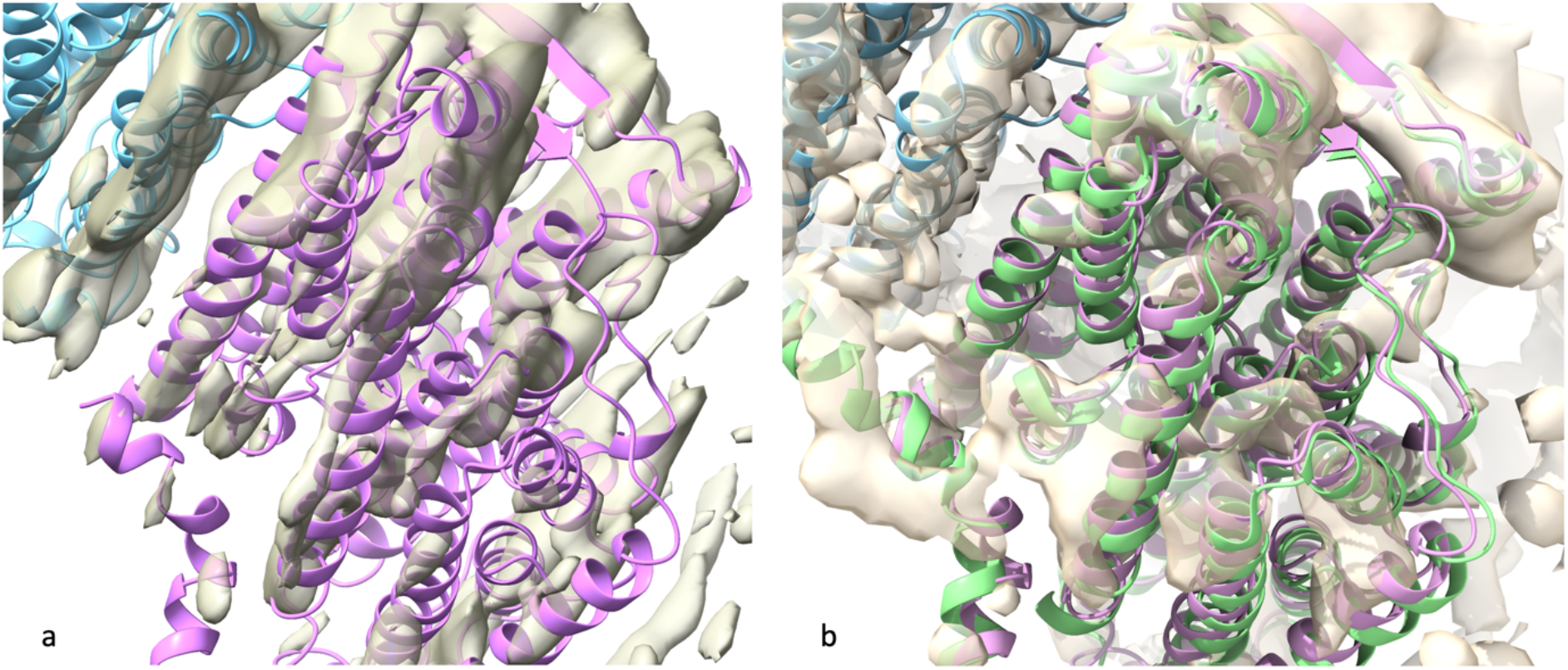
Model derived from chain L (magenta) of PDB entry 3rko, docked into the region of the map corresponding to chain L of PDB entry 7nyu (EMDB entry EMD-12654). Chain M of PDB entry 7nyu is shown in light blue. a) The map is computed using the Fourier coefficients 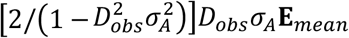 arising from the analysis of the local map volume (Millán et al., 2023). b) Chain L of PDB entry 7nyu is shown in light green, and the map is the full map from EMDB entry EMD-12654.

#### 4.1.2 AlphaFold model of megabody bound to GABA receptor

The structure of the pentameric human *γ*-aminobutyric acid (GABA) receptor bound to a megabody (PDB entry 7a5v, EMDB entry EMD-11657) was determined at a high overall resolution of 1.7 Å (Nakane *et al*., 2020), and models of the GABA receptor monomers are placed easily. However, the local order of each megabody bound to the periphery of the pentamer is significantly worse than that of each GABA receptor monomer, dropping to a local resolution of about 3 Å at the periphery as judged by *phenix*.*local_resolution*. To prepare a search model for the megabody (for which there is no independent structure), its structure was predicted using the community ColabFold version (Mirdita *et al*., 2022) of AlphaFold (Jumper *et al*., 2021), and the model was processed through *phenix*.*process_predicted_model* to remove very low-confidence residues and down-weight the lower-confidence remaining residues (Oeffner *et al*., 2022). The overall reconstruction has 5-fold symmetry, but only two of the five copies were placed in the global search. We note that this search took more than 30 minutes, longer than any of the other global search test cases (Millán *et al*., 2023).

We wished to understand why the global search failed to find all five copies and to explore whether *emplace_local* would be an effective alternative to the global *em_placement* search in similar situations. In this particular case, missing copies can easily be generated by applying the 5-fold symmetry to one or other of the two placements, but in the general case, the symmetry may be inexact or unknown. Local searches centered at each of the five megabody positions gave similar results (Table 2): each search took about 55 seconds, correctly placing a copy of the megabody with an LLG score above 500. In each case, the correct orientation was the top hit in the rotation search, of around 6 trial orientations. When run from the ChimeraX interface with the symmetry option enabled, the solutions were correctly expanded over the 5-fold symmetry.

**Table 2.**
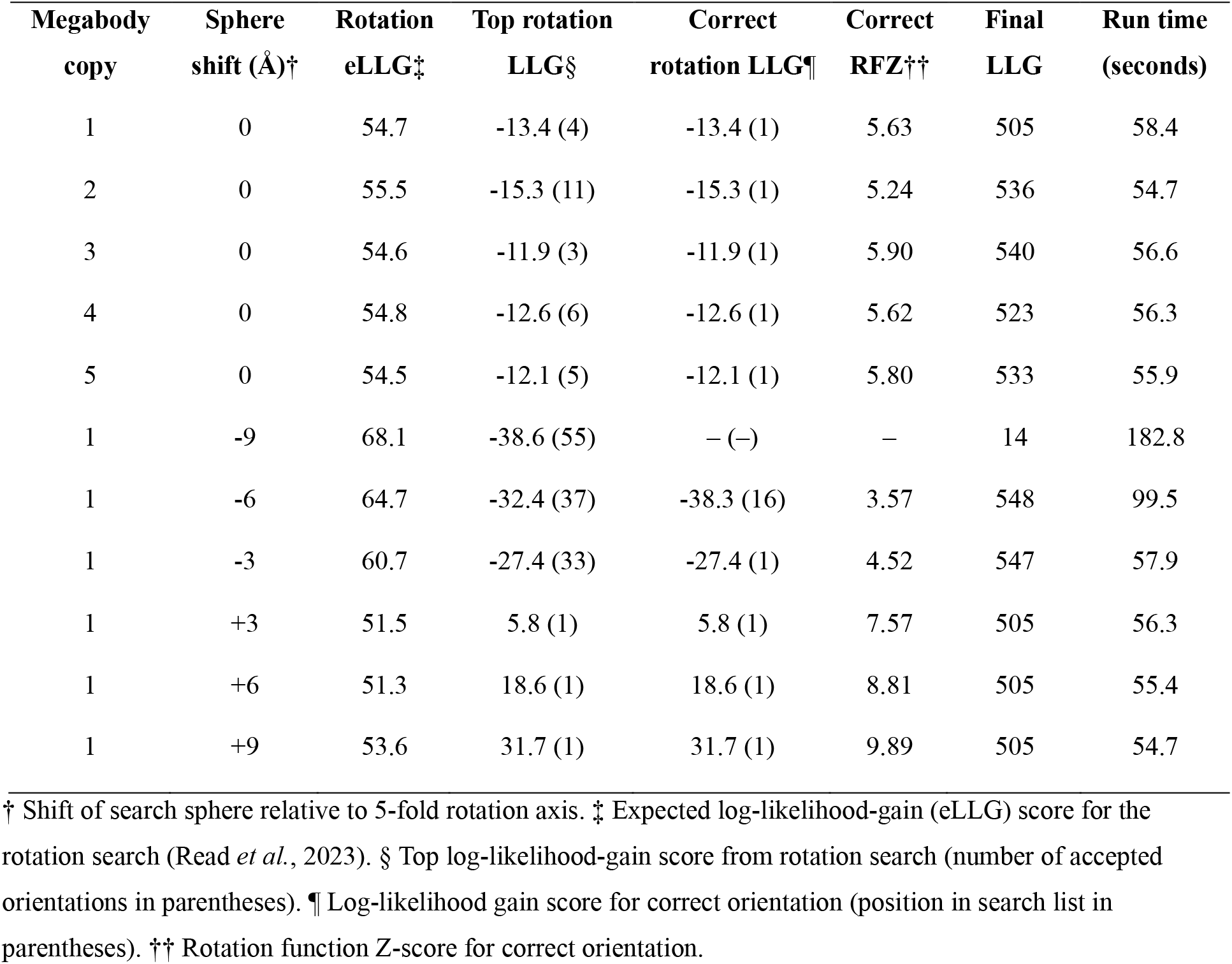
Results of *emplace_local* searches for megabody in complex with GABA receptor.

Varying the center of the search sphere had a substantial effect on the efficiency of the search. Since the megabody on the periphery is less well-ordered than the GABA receptor in the center of the reconstruction, it seemed reasonable to consider that varying the search center could alter the degree to which the GABA receptor component contributed to the search volume, adding strong signal that could not be explained by the megabody model and thus contributing to noise in the search. Moreover, when noise from the unexplained signal extends to higher resolution than the target signal, the likelihood target is mis-calibrated to expect more signal and less noise at high resolution. This reasoning accounts very well for what was observed when the search sphere for the first copy of the megabody was systematically moved nearer and farther from the 5-fold symmetry axis, thereby including more or less of the GABA receptor density respectively. The search sphere was moved in 3 Å steps from 9 Å nearer to 9 Å farther. With a sphere radius of 25.7 Å, two spheres offset by as much as 9 Å still share 74% of the same volume. Nonetheless, when the search sphere was shifted 9 Å closer to the rotation axis (−9 Å), the search failed because the correct orientation was not found in the default rotation search. When the search sphere was moved farther from the rotation axis, the search succeeded and the statistics (clarity of rotation search, run time) continued to improve as the search sphere was shifted outward to include less overlap with the GABA receptor region of the map (Table 2). The expected log-likelihood-gain (eLLG) score increases as the search sphere moves closer to the rotation axis, because there is more ordered density to contribute to the signal that is detected in the eLLG calculation. However, because the most well-ordered density largely arises from parts of the map that are not explained by the megabody model, it actually contributes additional noise to the search, leading to lower and even negative rotation LLG values for the spheres closest to the rotation axis. Note that negative LLG values for both rotation and translation searches arise when the model predicts the data less well than expected from the presumed quality of model and data. The variation in the final LLG score for the successful searches for the first copy of the megabody was unexpected but arises from a stochastic difference of one grid spacing in the centers chosen for the spheres used for the final rigid body refinement; the lower scores correspond to grid centers slightly nearer the 5-fold axis.

The sensitivity of the rotation search to the presence of more strongly-ordered density from the GABA component explains the limited success of the global search. In the global search, the reconstruction was covered by 6 overlapping spheres with radii of 58.2 Å, which did not follow the 5-fold symmetry. These spheres are likely to overlap significantly with the GABA component. In addition, because the expected LLG for the rotation search scales inversely with sphere volume (Read *et al*., 2023), only about 1/12 of the signal expected for the local search sphere would be expected in the subvolumes for the global search.

#### 4.1.3 Constant region of F_ab_ bound to α3β4 ganglionic nicotinic receptor

The structure of the α3β4 ganglionic nicotinic receptor bound to the ligand AT-1001 and to the F_ab_ fragment from a monoclonal antibody (PDB entry 6pv8, EMDB entry EMD-20488), was determined by cryo-EM at a resolution of 3.87 Å (Gharpure *et al*., 2019). Copies of the F_ab_ fragment bind to the two α3 subunits in the heteropentamer. In the published structure, only the variable domains directly contacting the receptor were fit, because the map quality is poor for the constant domains (Fig. 3a). As judged using *phenix*.*local_resolution*, the local resolution is about 5.0-5.5 Å in this region. Unfortunately, the box chosen for the map deposition has a boundary very near to the constant domains. This makes it difficult to extract appropriately-sized spheres of density for the docking algorithm; the map analysis assumes that the solvent region has the same noise distribution as the protein region, so it is not appropriate to pad the input map with zeros.

**Figure 3.**
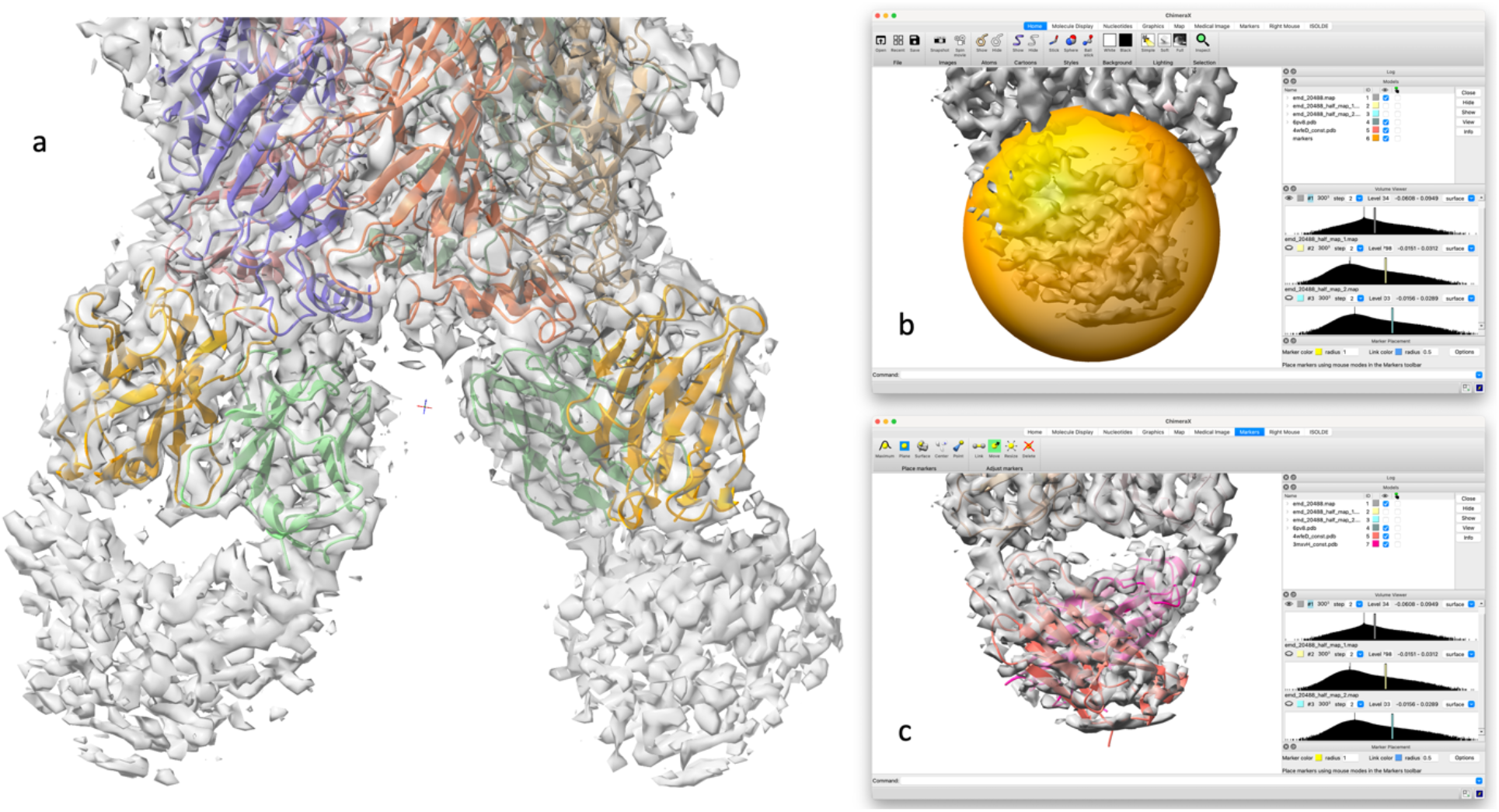
Searching for the F_ab_ constant domains in the complex with the α3β4 ganglionic nicotinic receptor (EMD-20488). a) The published structure is shown in the deposited map. The modelled part of the F_ab_ fragments consists of the variable domains of the heavy chain (light orange) and the variable domains of the light chain (light green). The map is of much poorer quality for the constant domains of these chains, at the bottom of the image. b) The search sphere, corresponding to the uninterpreted map region at the lower right of panel (a), is indicated by the yellow ball. c) The placed model for the constant domain from chain D of PDB entry 4wfe is shown as an orange cartoon, while the model derived from chain H of 3mxv is shown in pink.

We wished to see whether models of the constant domains could be docked, either with the global *em_placement* search or with *emplace_local*. In the published structure, the initial model for the variable domain of the light chain was derived from PDB entry 4wfe (Brohawn *et al*., 2014), and the variable domain of the heavy chain from PDB entry 3mxv (Maun *et al*., 2010). Our models of the corresponding constant domains were derived from the same PDB entries. Although there are two independent copies of the F_ab_ bound, only the one interacting with the α3 subunit in chain D of 6pv8 is sufficiently far from the edge of the deposited reconstruction.

Global docking trials of the constant domains with *em_placement* failed for reasons similar to those encountered in the previous two test cases: strong noise from better-ordered parts of the map that contained other components misled the search algorithm. When docking the constant domain from the light-chain model (chain D of 4wfe) in the global search, three potential solutions were found with LLG values varying from 35 to 70. The solutions are non-random, but they incorrectly superimpose beta-structure from the constant domain on beta-structure from the better-ordered components. Docking the constant domain from the heavy-chain model (derived from chain H of 3mxv) gave similar incorrect results, with LLG values varying from 23 to 58 for five potential solutions, which are all incorrect but non-random.

In contrast, local docking with *emplace_local* succeeded when care was taken to avoid searching in the part of the map occupied by the variable domains. This was easiest to achieve with the interactive ChimeraX interface. Fig. 3b shows the positioning of the search sphere in ChimeraX to avoid the part of the map accounted for by the variable domains of the relevant copy of the F_ab_ (interacting with the α3 subunit in chain D of 6pv8), while Fig. 3c shows the result of searching within this volume for the two constant domain models. The light-chain constant domain model gave an LLG score of 159 and a map correlation of 0.473, while the heavy-chain constant domain model yielded LLG=135 and a map correlation of 0.481.

### 4.2. Local searches without half-maps

Half-maps are required in *em_placement* to estimate the direction- and resolution-dependence of the signal and noise in the likelihood target. The approximation that we use in *emplace_local* when half-maps are unavailable ignores the direction dependence and assumes that the resolution dependence of signal-to-noise can be modelled by a simple isotropic exponential falloff governed by the estimated local resolution of the map, given in (1). These approximations might be expected to limit the success of this method, especially when the local signal is poor or the appropriate local resolution is uncertain. For this reason, we chose to look in detail at test cases for which the global docking search failed, and where signal-to-noise was marginal even for local docking searches.

#### 4.2.1 Chain L of the *E. coli* respiratory complex I

As noted above, the local map resolution around chain L is estimated by the original authors to be about 9-11 Å, although the nominal (best) resolution for the whole reconstruction is 3.8 Å. We ran a series of local docking tests using either half-maps or the full map, and varying the nominal resolution limit (Table 3). If half-maps were used, the docking searches were successful for all resolution limits tested, from 3.8-13 Å. However, when the full map was used the docking searches failed for resolution limits better than 7 Å (*i*.*e*., when the resolution was indicated to be better than it actually is), confirming that the method using a single full map is indeed more sensitive to the choice of resolution limit. The full-map docking calculation at higher resolution limits was drawn into fitting the better-ordered map features of chain M.

**Table 3.** Effect of resolution on docking searches with half- and full-maps

| $d_{\min}$ (Å) | Complex I chain L | | |
| --- | --- | --- | --- |
|  | Top LLG<sup>†</sup> | Top incorrect | Run time (seconds) |
| Half-maps |  |  |  |
| 3.8 | 49.9 | — | 33.4 |
| 5.0 | 58.8 | — | 25.3 |
| 7.0 | 90.1 | — | 17.8 |
| 9.0 | 88.3 | — | 16.0 |
| 11.0 | 78.3 | — | 15.6 |
| 13.0 | 48.6 | — | 15.7 |
| Full map |  |  |  |
| 3.8 | 104.5<sup>‡</sup> | 104.5 | 122.8 |
| 5.0 | 116.9<sup>‡</sup> | 116.9 | 67.7 |
| 7.0 | 106.5 | — | 13.7 |
| 9.0 | 83.6 | — | 10.2 |
| 11.0 | 66.3 | — | 10.5 |
| 13.0 | 53.3 | — | 10.5 |
| Megabody bound to GABA receptor |  |  |  |
| $d_{\min}$ (Å) | Top LLG | Top incorrect | Run time (seconds) |
|  | Half-maps |  |  |
| 1.7 | 505.4 | — | 57.8 |
| 3.0 | 608.0 | — | 36.3 |
| 5.0 | 367.3 | — | 27.8 |
| 7.0 | 218.6 | — | 27.8 |
| 9.0 | 59.2<sup>‡</sup> | 59.2 | 36.1 |
| 11.0 | 47.9<sup>‡</sup> | 47.9 | 27.1 |
| Full map |  |  |  |
| 1.7 | 629.9 | — | 35.0 |
| 3.0 | 541.0 | — | 19.9 |
| 5.0 | 211.8 | — | 14.6 |
| 7.0 | 106.2 | — | 14.3 |
| 9.0 | 66.1 | 45.4 | 18.0 |
| 11.0 | 27.6<sup>‡</sup> | 27.6 | 13.7 |
| F<sub>ab</sub>-H constant domain |  |  |  |
| $d_{\min}$ (Å) | Top LLG | Top incorrect | Run time (seconds) |
|  | Half-maps |  |  |
| 3.87 | 118.0 | — | 20.3 |
| 5.0 | 158.9 | — | 19.9 |
| 7.0 | 150.1 | — | 20.4 |
| 9.0 | 140.3 | — | 19.1 |
| 11.0 | 85.8 | 40.3 | 29.2 |
| 13.0 | 62.9 | — | 18.1 |
| Full map |  |  |  |
| 3.87 | 156.1 | — | 11.1 |
| 5.0 | 147.0 | — | 10.5 |
| 7.0 | 91.6 | — | 10.8 |
| 9.0 | 61.1 | — | 10.1 |
| 11.0 | 47.3 | 34.5 | 16.5 |
| 13.0 | 34.5 | 26.6 | 11.3 |

<sup>†</sup> Top log-likelihood-gain score from docking search. <sup>‡</sup> Incorrect solution.

Ideally, a proper calibration of the error model in the likelihood target, based on the statistical analysis of half-maps, should account for the lack of information in higher resolution Fourier terms than the local resolution, effectively ignoring them. However, we see that the inclusion of high-resolution data degrades the performance of the docking algorithm in this case, even when half-maps are used, indicating that there is still room for improvement in the algorithms estimating signal and noise contributions.

#### 4.2.2. AlphaFold model of megabody bound to GABA receptor

In this case, the imposition of 5-fold symmetry on the reconstruction has helped to ensure that there is little anisotropy in the signal or noise. As a result, the approximation used for the full-map searches is more appropriate than in the other test cases, and the success rates are very similar (Table 3). In fact, searches imposing a resolution limit of 9 Å fail with the half maps but succeed with the full map, although the separation between the correct score and the top incorrect score is relatively small.

#### 4.2.3 Constant region of F_ab_ bound to α3β4 ganglionic nicotinic receptor

This test case is complicated by the fact that the deposited reconstruction has been truncated close to the unmodelled domains. The heavy-chain constant domain from the F_ab_ bound to chain D is furthest from the map boundary, so the model derived from chain H of 3mxv was used for the full map docking comparisons.

Both sets of searches, using half-maps or full maps, succeed over the full range of resolution limits tested, from 3.87-13.0 Å. However, the half-map searches yield less ambiguous answers and, although the full-map search at 13.0 Å resolution succeeds, there is very little distinction between the correct placement and the top incorrect solution.

Note that, for all test cases, the single-map protocol is up to twice as fast when it succeeds, because it does not require the likelihood-based estimation of signal and noise parameters. In addition, in a variety of tests for cases with better signal strength, the single-map protocol was found to be robust and efficient (results not shown).

### 4.3 Comparison with ChimeraX *fitmap* algorithm

The Local EM Fitting procedure complements an existing *fitmap* algorithm integrated into ChimeraX. Most commonly, *fitmap* is used to optimize the rigid-body fit of a model that has already been placed approximately correctly, but it can optionally be used to test a number of random orientations and positions around a selected location in the map.

We tested *fitmap* on the test cases used for *emplace_local*, initially using default settings for the random local search where possible. To restrain the search to a local region, the *radius* keyword must be given, and we used a value of half the radius of a sphere enclosing the search model to make the search volume comparable to what is covered by *emplace_local*. The search is carried out around the starting position of the model, which was placed for these tests by superimposing it on the deposited structure. (Control tests with the position offset by a fraction of the model radius gave similar results, not shown.) The number of random trials (*search* keyword) was set initially to 1000, which made the computing requirements comparable to those of *emplace_local*. The default score that is optimized, for fitting a model in a map, is the average map value at atomic positions in the model. If the *resolution* keyword is supplied, then a map-to-map search is performed, scored by the integral of the product between a map generated at that resolution from the placed model and the experimental map. The results of both search methods vary with the level of smoothing or sharpening applied to the input map, in contrast to the *emplace_local* method where the effects of noise at high resolution are accounted for by the variation in Fourier space of the noise power estimates. Different levels of smoothing were not explored in this work. For *fitmap*, the *resolution* parameter should preferably be set to the local map resolution, but a variety of values was tested to determine the sensitivity to this parameter.

#### 4.3.1 Chain L of the *E. coli* respiratory complex I

Using the command

~~~
fitmap #2 in #1 search 1000 radius 20.5
~~~

where the trimmed model from chain L of PDB entry 3rkoL was display item 2 and the deposited map was display item 1, the correct placement (with an average map value score of 1.444) was found in 10 of the 1000 trials, taking a total of 44.6 seconds. If either the search model was moved closer to the position of chain M in the deposited structure or the search radius was increased to 80 Å, the *fitmap* algorithm was led to find an incorrect superposition on chain M with the higher score of 1.796, similar to the single full-map search with *emplace_local* when run at resolutions higher than 7 Å.

The *fitmap* procedure in ChimeraX also succeeded when a *resolution* keyword was given to activate the map-to-map fit, for a range of resolutions between 3.8 and 9.0 Å. Run times were considerably longer than the comparable runs using the *emplace_local* procedure (Table 3), taking from 46.2 seconds at 9.0 Å to 186 seconds at 3.8 Å. When a resolution of 11.0 Å was chosen for *fitmap*, the correct placement was second in the list of possibilities, with a score of 0.599 compared to 0.602 for the highest incorrect result. By comparison, *emplace_local* succeeded at resolutions as low as 13.0 Å, using either the full map or half-maps (Table 3).

#### 4.3.2 AlphaFold model of megabody bound to GABA receptor

Three sets of 1000 search trials were carried out using the *fitmap* command with a radius of 24.0 Å, each taking about 43 seconds. In one of these sets, the correct placement with an average map value score of 0.027 was found twice, but in the other two sets of trials the search failed, yielding a top score of 0.015. A larger-scale search found the correct solution in 5 of 5000 trials, taking 211 seconds. To be confident of finding the correct solution in the random search for this problem, we conclude that several thousand trials would indeed be required. By comparison, the successful searches using the deterministic *emplace_local* algorithm took about 55 seconds (Table 2).

The map-to-map option was somewhat more successful, giving 3-8 solutions per 1000 trials when the resolution was specified in the range of 1.7-5.5 Å (taking 85.2 to 157 seconds), but failing or yielding only a single solution when the resolution was set lower. In contrast, *emplace_local* succeeded when using half-maps with the resolution specified between 1.7 and 7.0 Å and with just the single full map at resolutions as low as 9.0 Å, with each of these calculations taking less than a minute (Table 3).

#### 4.3.3. Constant region of F_ab_ bound to α3β4 ganglionic nicotinic receptor

The ChimeraX *fitmap* command was tested for docking the same copy of the heavy chain constant domain (derived from chain H of PDB entry 3mxv) as the full-map *emplace_local* tests. A default search with 1000 trials and a radius of 22.7 Å ran in 34.4 seconds; the correct solution was found 8 times with a score of 0.00805, but an incorrect solution (superimposed on the light chain) had the slightly higher score of 0.00820. The map-in-map option was again more successful, with the top-scoring solution being correct for resolutions of 3.87-9.0 Å, taking from 35 to 58 seconds. With a resolution of 11.0 Å, the correct solution (found 5 times) was third in the list with a score of 0.734, compared to 0.756 for the top solution. At 13.0 Å, the correct solution was not found, even with as many as 10,000 trials.

## 5 Discussion and conclusions

Generally, it is preferable to carry out an unbiased search over a full cryo-EM map for a component of a complex or multi-domain protein. However, there are common situations in which an unbiased search is unsuccessful but there are good hypotheses for the location of a component. The *emplace_local* program was built using a subset of features of the global *em_placement* program, with the expectation that a local search would be faster and could be more sensitive than a global search. The results presented here demonstrate that the expectation was well-founded.

The local search typically takes under a minute to run, in the range required for a useful interactive tool. A new plugin for the interactive molecular graphics program, ChimeraX (Goddard *et al*., 2018), has therefore been developed, as part of a project to incorporate algorithms from the Phenix package (Liebschner *et al*., 2019) into ChimeraX.

The local docking approach relaxes the strict requirement for half-maps, which is very useful for legacy cryo-EM structures. The single-map protocol is somewhat less sensitive than the protocol using two half-maps, and it is necessary to specify the local map resolution (derived, for instance, from ResMap (Kucukelbir *et al*., 2014) or *phenix*.*local_resolution*), but it can work well in a variety of cases. In many cases, the *fitmap* global fitting procedure built into ChimeraX also works, but the *emplace_local* approach has several advantages. First, it gives log-likelihood-gain scores on an absolute scale that are independent of any scaling or overall sharpening applied to the maps; the user can be confident that scores around 60 are plausible, and scores much higher are very likely to be correct. In contrast, the *fitmap* scores depend on the mode chosen, with the absolute values depending on scaling and optimal results depending on the choice of sharpening parameters for the map. Second, *emplace_local* is deterministic, so that there is no concern that a larger number of trials might have found a solution that was randomly missed. Third, there is no need to switch modes for different cases, depending on factors like the resolution of the map; what matters is that the model is large enough to explain significant variations within the map or maps.

There are substantial advantages to having the graphical ChimeraX interface. Some of the most difficult docking problems involve placing poorly-ordered components next to components that are more well-ordered. Including well-ordered non-target regions of the map in the search volume adds noise to the search, but being able to visualize the search volume makes this easier to avoid. In addition, the user gains rapid visual feedback on the plausibility of the docking result, to complement the numerical likelihood and map correlation scores. An alternative and potentially powerful approach to handling noise from well-ordered neighbouring components would be to account for what has already been learned from placing models in that region, and this will be explored for future versions of *em_placement* and *emplace_local*.

## 6 Funding information

The following funding is acknowledged: Wellcome Trust (grant No. 209407/Z/17/Z to Randy J. Read); National Institutes of Health, National Institute of General Medical Sciences (grant No. P01GM063210 to Paul D. Adams, Randy J. Read and Thomas C. Terwilliger, and grant No. R24GM141254 to Paul D. Adams, which funds the ChimeraX collaboration). This work was supported in part by the US Department of Energy under Contract DE-AC02-05CH11231.

## Acknowledgements

We thank Tom Goddard for advice on the ChimeraX interface and for valuable comments on the manuscript.

## Notes

### Competing Interest Statement

The authors have declared no competing interest.

